# Discovering genetic loci associated with rate of vegetative index gain using UAV-based phenomics in spring wheat

**DOI:** 10.64898/2026.05.22.727186

**Authors:** Sana Ur Rehman, Ali Raza, Zijian He, Li Lei, Zahid Mahmood, Muhammad Fayyaz, Muhammad Waqas, Muhammad Salman Akhtar, Jiajie Wu, Yonggui Xiao, Muhammad Adeel Hassan, Zhonghu He, Awais Rasheed

## Abstract

In wheat, the pre-heading stage determines spikelet formation, floret fertility, and canopy development, making it a critical window for early stress detection and yield potential. The genetic basis of pre-heading canopy development in wheat has remained constrained by the conventional phenotyping due to the low temporal resolution. Here, we quantified the rate of vegetation index gain (RVIs) during tillering to heading stages using UAV-mounted multispectral sensor in 196 spring wheat cultivars representing 112 years of breeding history. RVIs were calculated using six vegetation indices for consecutive two growing seasons, and genome wide association study (GWAS) was performed on RVIs, grain yield (GY) and thousand grain weight (TGW) using a wheat 37K SNP array. RVIs showed significant positive correlations with grain yield (r=0.28-0.43; p<0.001) and consistently increased in the modern cultivars compared to old cultivars. This indicated that resource remobilization during pre-heading canopy development significantly contributed to GY during modern wheat breeding. GWAS identified 67 loci, including 12 Group-I loci associated only with RVIs, and 18 Group-II loci associated with both RVIs and yield traits. Two stable loci on chr1B and chr5D consistently increased GY and RVIs across environments, and the tag SNPs were converted to selectable KASP markers. The allelic distribution on global wheat collection of ∼3000 accessions showcased that favorable alleles on both loci were dominant in cultivars compared to landraces. Similarly, favorable alleles showed more frequency in winter type than spring type. Across breeding eras both alleles showed increasing trend with chr5D reaching near fixation and chr1B remaining partially enriched in modern cultivars. Our work on capturing pre-heading canopy development, discovery of two stable loci underpinning yield and RVIs, and development of KASP markers provided a strong foundation to HTP assisted genetic dissection of GY and facilitated the understanding of canopy dynamics and yield formation.

## 1. Introduction

Wheat (*Triticum aestivum* L.) is the most widely cultivated crop globally, contributing approximately 18% of the calories and 19% protein consumed by humans (Shewry, 2024). With the global population projected to exceed 9.7 billion by 2050, annual productivity gains of 1.6–2.4% are required to meet demand under increasing climate challenges (Ray et al., 2013). Addressing this challenge requires the discovery of novel physiological traits (Reynolds et al., 2022) and their genetic basis using modern phenotyping methods could help in capturing the missing heritability of quantitative traits like GY (Farooq et al., 2024; Rasheed et al 2025). GY in wheat is a complex trait conventionally measured by spike number per unit area, grains per spike, and grain weight (Calderini et al., 2021).

The Green Revolution in mid 1960s increased GY, mainly driven by increasing grain number per unit area, which is largely determined during the tillering to heading period. During the pre-heading stage, rapid canopy expansion, efficient radiation interception, and extended green-area duration enhance carbon assimilation, spike fertility, and carbon allocation, ultimately determining biomass accumulation and grain number (Foulkes et al., 2011, 2022; Reynolds & Braun, 2022). Within the source-sink framework, green biomass serves as the primary source of assimilates, while developing spikes act as the main sinks. Genotypes with greater green biomass accumulation or more efficient coordination between biomass production and sink demand generally achieve higher yield potential (Murchie et al., 2023). Therefore, phenotyping during the vegetative stage is critical, as it also captures early stress responses that can directly influence biomass accumulation and spike formation; limitations at this stage cannot be fully recovered during reproductive stage, making it a key window for identifying genotypes with superior growth and stress resilience that underpin high yield (Çakir, 2004).

However, quantifying these physiological processes under field conditions has been challenging due to reliance on destructive and low-throughput methods, leading to the “phenotyping bottleneck” (Fiorani & Schurr, 2013). In recent years, non-destructive, imaging-based high-throughput phenotyping (HTP) platforms ranging from ground-based systems to UAVs and satellite remote sensing have been developed to address this bottleneck. These HTP approaches facilitate the quantification of relationships between plant growth processes and yield formation (Gao et al., 2023; Mir et al., 2019). Among these platforms, UAVs equipped with multiple sensors such as RGB, multispectral, hyperspectral infrared thermal imagers, and LiDAR have emerged as practical solution for field-based phenotyping (Yang et al., 2017). UAVs enable rapid imaging of thousands of breeding plots at centimeter-scale resolution across the entire growing season, generating dense time-series data with high heritability for canopy development, biomass accumulation, and stress responses (Tattaris et al., 2016; Xie & Yang, 2020).

In wheat, UAV-derived spectral vegetation indices (VIs) have been widely used as non-destructive quantitative proxies for evaluating a range of agronomic and physiological traits, including plant height (Hassan et al., 2019), chlorophyll content (Cheng et al., 2025), nitrogen status (Yang et al., 2020), biomass (Zhao et al., 2025), leaf area index (Mulero et al., 2025), and senescence dynamics (Hassan et al., 2018). Beyond serving as proxies for static agronomic traits, time-series VIs collected across the growing season further enable digital-agriculture applications, such as crop modeling, machine learning, deep learning and genomic selection (Farooq et al., 2024). Studies utilizing these VIs achieved substantially increased accuracy in predicting green biomass and GY. For instance, Fei et al., (2021) achieved an R^2^ of 0.73 for yield prediction by combining UAV based multispectral and thermal infrared data. More recent work has demonstrated that integrating multi-stage spectral features with additional canopy trait parameters can improve prediction performance for yield with accuracy ranging R^2^≈0.85–0.95 under optimal measurement stages and modeling frameworks (Bian et al., 2022; Ge et al., 2025; Kang et al., 2024; Zhou et al., 2023). UAV-derived traits can serve as effective secondary covariates in genomic selection, enhancing prediction accuracy (Crain et al. 2018; Kaushal et al. 2024). These findings indicate that UAV-based VIs can provide robust and cost-efficient proxies for key physiological processes such as radiation-use efficiency, biomass accumulation, and stress responses. However, despite their increasing use in phenotyping and genomic prediction in wheat, the use of UAV-derived VIs for identifying genetic loci underlying these traits remains limited.

GWAS is a powerful tool widely used to dissect the genetic architecture of complex quantitative traits by leveraging historical recombination and allelic diversity across diverse wheat germplasm panels. However, the statistical power, mapping resolution, and biological interpretability of GWAS depend strongly on the precision and temporal richness of phenotypic measurements (Yang et al., 2014). This limitation becomes particularly critical for highly polygenic, quantitative traits such as GY, where genetic effects are small, developmentally regulated, and strongly modulated by genotype-by-environment (G×E) interactions (Rasheed et al., 2025). Several studies have used UAV-derived traits-including plant height dynamics, responses to salinity and waterlogging, senescence progression, and canopy cover for quantitative genetic analyses (Hassan et al., 2021; Kang et al., 2024; Singh et al., 2019). However, only a limited number of studies in wheat have incorporated multi-temporal UAV-derived VIs into genetic analyses, with most focusing on late-season senescence traits (Hassan et al., 2021; L. Li et al., 2023). To date, no study has integrated GY with the temporal dynamics of VI accumulation coupled with genetic studies during the tillering-to-heading phase, which is critical period for yield determination in wheat. Therefore, the aims of this study are to, (1) quantify the temporal rate of increase in multispectral VIs during pre-heading window using high-resolution UAV time-series data (2) conduct GWAS integrating GY, TGW and pre-heading vegetative index gain in a diverse panel spring wheat genotypes and (3) identify and characterize stable loci controlling the rate of pre-heading vegetative index gain and its genetic relationship with GY. We further surveyed the global wheat germplasm collection to identify the distribution of favorable alleles at two stable loci.

## 2. Material Method

### 2.1 Germplasm and field trial

A natural population consisting of 196 historical and modern bread wheat genotypes released in Pakistan between 1911 and 2023 was evaluated over two consecutive growing seasons, 2022–2023 (E1) and 2023–2024 (E2), at the National Agricultural Research Centre (NARC), Islamabad, Pakistan (33.67°N, 73.13°E). To optimize UAV-based high-throughput phenotyping and reduce inter-plot spectral interference, the experiment was arranged in a non-contiguous (skip-plot) design, where each planted plot (5 m × 1.5 m; six rows with 25 cm spacing) was separated by unplanted rows in both directions, forming a checkerboard layout. Standard irrigation and agronomic practices for spring wheat recommended for the region were applied uniformly across all plots.

### 2.2 UAV-based phenotyping

#### 2.2.1 Data acquisition and processing

High-throughput phenotyping was performed using a DJI Inspire 2 drone equipped with a Sentera Double 4K multispectral sensor (12.3 MP, five bands: Blue (B), Green(G), Red (R), Red Edge (RE), Near-infrared (NIR) (Sentera, 2024). In each season E1 and E2, ten flights were conducted at 30 m above-ground altitude with 80% frontal and 75% lateral overlap, exclusively covering the tillering to heading period (Zadok’s GS20–GS59). All flight missions were pre-programmed and autonomously executed using the DJI Pilot 2 app to ensure reproducible flight paths and imaging geometry. Flights were performed under clear sky conditions between 11:00 and 13:00 local time to minimize bidirectional reflectance effects. Raw images were processed in Pix4Dmapper (Pix4D, 2025) using the Ag Multispectral template, producing georeferenced Ortho mosaics (ground sampling distance ≈ 1.2 cm pixel⁻¹) in WGS84 / UTM zone 43N projection (Figure 1).

**Figure 1.**
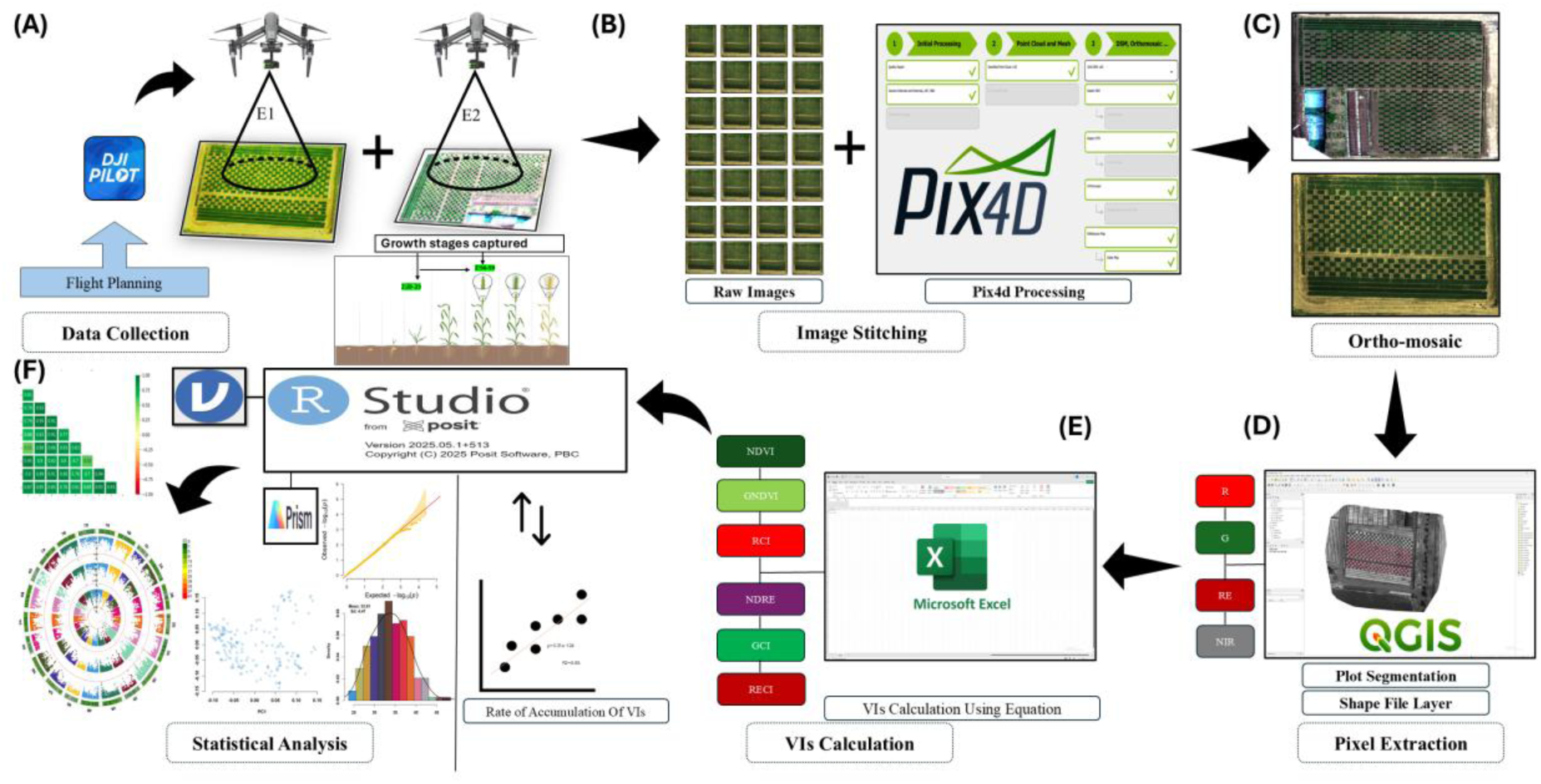
Workflow of the high-throughput phenotyping and GWAS pipeline, **(A)** Multi-temporal UAV multispectral data acquisition, **(B)** Image stitching and radiometric calibration, **(C)** Ortho-mosaic formed after stitching, **(D)** Plot-level segmentation, individual band extraction (R,B,G, RE,NIR), **(E)** Calculation of VIs and rate-of-Gain, **(F)** GWAS and other statistical analysis.

#### 2.2.2 Plot segmentation and VIs calculation

Orthomosaic images were processed in QGIS v3.44.0 (Dawson et al., 2025) for plot segmentation (Figure 1). Plot boundaries were defined using shapefile layers, and each polygon was assigned to a corresponding genotype ID (n = 196). Prior to reflectance extraction, non-canopy pixels (soil and background) were excluded using NDVI-based masking to retain only vegetation signals. Mean reflectance values for each plot were then extracted using the Zonal Statistics tool in QGIS and exported to Microsoft Excel for VI computation based on the equations provided in (Table 1). Subsequently, six VIs were used to estimate their rates of accumulation for each plot from tillering (Zadok’s GS20) to heading (Zadok’s GS59).

**Table 1.** Vegetation indices derived from UAV-based multispectral imagery, including their formulas and applications in phenotyping.

| Indices | Equation | Application | Reference |
| --- | --- | --- | --- |
| NDVI (Normalized Difference Vegetation Index) | $\frac{(NIR - R)}{(NIR + R)}$ | General plant health, biomass, yield. | (Rouse et al., 1974) |
| GNDVI (Green NDVI) | $\frac{(NIR - G)}{(NIR + G)}$ | Chlorophyll, nitrogen status, yield, plant vigor. | (Gitelson et al., 1996) |
| NDRE (Normalized Difference Red Edge) | $\frac{(NIR - RE)}{(NIR + RE)}$ | Chlorophyll, nitrogen, stress detection. | (Gitelson & Merzlyak, 1994) |
| GCI (Green Chlorophyll Index) | $\frac{(NIR)}{(G - 1)}$ | Chlorophyll content, yield. | (Gitelson et al., 1996) |
| RCI (Red Chlorophyll Index) | $\frac{(NIR)}{(R - 1)}$ | Pigment status via red absorption (sensitive to stress onset). | (González-Espíndola et al., 2024) |
| RECI (Red Edge<br>Chlorophyll Index) | $\frac{(NIR)}{(RE - 1)}$ | Canopy<br>chlorophyll<br>content, N<br>status. | (Gitelson et al.,<br>2005) |

#### 2.2.3 Calculation of rate of vegetation index (RVIs) gain

The rate of accumulation for each vegetation index (RVIs) was computed as the average change in VI between consecutive flights, standardized by the number of days between flights. For each plot this was expressed as:

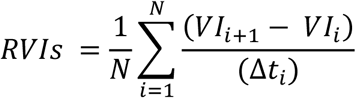

Here, RVIs represents rate of accumulation of VIs, 𝑖 denotes each consecutive flight interval, N is the total number of such intervals, 𝑉𝐼_𝑖_ and 𝑉𝐼_𝑖+1_ are VI values from flights I and 𝑖+1, and Δ𝑡_𝑖_ is the corresponding number of days between flights.

### 2.3 Grain yield and thousand grain weight

At physiological maturity, GY and TGW were collected manually using standard field protocols that align with CIMMYT’s standard wheat phenotyping protocols (Reynolds et al., 2020). GY was measured by threshing and cleaning the harvested grain from each plot, then weighing it using a digital scale. The grain moisture content was assessed with a moisture meter, and yields were adjusted to a standard moisture level (typically 12–14%) to ensure accuracy. The adjusted GY was expressed in kg m^−2^. For TGW, a random sample of 1,000 fully developed, cleaned grains were counted manually, and weighed using a precision balance (0.01 g). The grain sample was dried to the standard moisture level before weighing, and TGW was recorded in grams.

### 2.4 Phenotypic data analysis

Statistical analyses were performed using R v4.4.3, Jamovi and GraphPad Prism 9. Principal Component analysis (PCA) was performed and visualized using R package FactoMineR. To account for environmental variation across the two field years, genotype Best Linear Unbiased Estimates (BLUEs) were derived from a linear model fitted using the lme4 package in R, with year and genotype treated as a fixed effect (Henderson, 1975; Piepho et al., 2007).

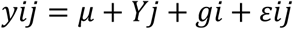

where 𝑦_𝑖𝑗_is the observed phenotypic value of the 𝑖^𝑡ℎ^genotype in the 𝑗^𝑡ℎ^year, 𝜇is the overall mean, 𝑌_𝑗_is the fixed effect of year, 𝑔𝑖is the fixed effect of genotype, and 𝜀_𝑖𝑗_is the residual error.

### 2.5 DNA extraction and genotyping

DNA was isolated from 20 mg fresh leaves using CTAB method described by (Dreisigacker et al., 2016). Quality of DNA was evaluated by electrophoresis on a 1% agarose gel, while concentration and purity were assessed using a NanoDrop spectrophotometer based on the A260/A280 absorbance ratio. The wheat cultivar panel was subsequently genotyped using the 37K SNP genotyping-by-targeted sequencing (GBTS) platform (wheat 16K Plant Comm), and data is publicly available (Rasheed & Fayyaz, 2023).

### 2.6 Genome-wide association analysis

Genotypic data from the 37K SNP array were subjected to quality control. SNPs with a minor allele frequency (MAF) < 5% were removed using TASSEL v5.2, resulting in 23,897 high-quality SNPs retained for GWAS. All subsequent GWAS analyses, including calculation of the kinship matrix (K) and population structure analysis (via principal component analysis, PCA), were performed in Rv4.4.3. A Q matrix was constructed from the first five principal components. GWAS were conducted using the rMVP package in R v4.3.3, implementing the Farm CPU model. Significant SNP-trait associations were declared at a threshold of −log10 (P) > 4.

### 2.7 Development of KASP marker

Kompetitive Allele-Specific PCR (KASP) assays were developed to validate the two target loci regions identified in the GWAS. The genomic sequences flanking the tag SNPs were retrieved from the Wheat Omics database (Ma et al., 2021). For each locus, two allele-specific forward primers were designed with standard FAM (5′-GAAGGTGACCAAGTTCATGCT-3′) and HEX (5′-GAAGGTCGGAGTCAACGGATT-3′) tails, with the target SNP positioned at the 3′ end, along with a locus-specific common reverse primer (Supplementary Table S1). Amplicon sizes were maintained below 120 bp to ensure efficient amplification. Primer mixtures were prepared by combining 46 µL ddH₂O, 30 µL common primer (100 µM), and 12 µL of each tailed allele-specific primer (100 µM). KASP reactions were performed in 96-well plates in a total volume of approximately 3 µL containing 10–20 ng genomic DNA, 1× KASP master mix, and 0.056 µL primer mix. PCR amplification was carried out with an initial hot start at 95 °C for 15 min, followed by ten touchdown cycles (95 °C for 20 s; annealing starting at 65 °C and decreasing by 1 °C per cycle for 25 s), and 30 additional cycles (95 °C for 10 s; 57 °C for 60 s). Fluorescence signals were recorded using a real-time PCR detection system, and genotype calls were assigned based on cluster separation (Rasheed et al., 2016).

## 3. Results

### 3.1 Phenotypic variations

Wheat varieties were grouped into three breeding eras to assess historical changes in phenotypes. First group include 10 pre-green revolution cultivars (Pre-GR), 38 cultivars released during green revolution (GR) i.e. 1965–1990, and 148 modern cultivars released after 1990. GY and TGW exhibited substantial variation across two environments E1 and E2 and their BLUEs. In E1, mean GY was 1.42 kg m⁻², ranging from 0.25 kg m⁻² to 2.50 kg m⁻². In contrast, E2 recorded a higher mean GY of 1.71 kg m⁻², ranging from 0.50 kg m⁻² to 2.75 kg m⁻². The average GY of Pre-GR were 1.2 kg m⁻² (E1), 1.2 kg m⁻² (E2), followed by GR 1.3 kg m⁻² (E1), 1.6 kg m⁻² (E2), and highest by modern cultivars 1.5 kg m^−2^(E1), 1.8 kg m^−2^ (E2). TGW exhibited moderate environmental variation, with a mean of 33.8 g in E1 (ranging from 23.8 to 46.2 g) and 36.3 g in E2 (ranging from 20.8 to 48.3 g).

The mean rate of RVIs gain during the tillering-to-heading stages varied substantially between environments and VIs. In E1, mean RVIs were slightly lower for most indices as compared to E2, ranging from lowest 0.0022 day⁻¹ (NDRE_E1) to highest 0.0632 day⁻¹ (RCI_E1). In contrast, E2 showed consistently higher mean RVIs, with RCI_E2 reaching 0.115 day⁻¹ and NDRE_E2 0.00371 day⁻¹ (Table 2).

**Table 2.** Summary of basic descriptive Statistic of phenotypic traits across both environments.

| Trait | Mean ± Standard Deviation | Range (min-max) | P-Value |
| --- | --- | --- | --- |
| GY_E1 | 1.42 ± 0.398 | 0.25 - 2.5 | <0.001 |
| GY_E2 | 1.71 ± 0.46 | 0.5 - 2.75 |  |
| TGW_E1 | 33.8 ± 4.47 | 23.8 - 48.3 | <0.001 |
| TGW_E2 | 36.3 ± 4.83 | 20.8 - 46.2 |  |
| GCI_E1 | 0.0271 ± 0.0104 | 0.00804 - 0.0588 | <0.001 |
| GCI_E2 | 0.00676 ± 0.00331 | 0.000865 - 0.0154 |  |
| GNDVI_E1 | 0.00336 ± 0.00187 | 0.0002 - 0.00948 | <0.001 |
| GNDVI_E2 | 0.0075 ± 0.00139 | 0.00303 - 0.0113 |  |
| NDRE_E1 | 0.00221 ± 0.00121 | 0.000062 - 0.0054 | <0.001 |
| NDRE_E2 | 0.00371 ± 0.000661 | 0.00181 - 0.00556 |  |
| NDVI_E1 | 0.00441 ± 0.00214 | 0.000249 - 0.0124 | <0.001 |
| NDVI_E2 | 0.0403 ± 0.00602 | 0.0267 - 0.0611 |  |
| RCI_E1 | 0.0632 ± 0.0265 | 0.00678 - 0.13 | <0.001 |
| RCI_E2 | 0.115 ± 0.019 | 0.075 - 0.199 |  |
| RECI_E1 | 0.00827 ± 0.00107 | 0.0053 - 0.0108 | <0.001 |
| RECI_E2 | 0.01 ± 0.0017 | 0.00531 - 0.0157 |  |

Modern cultivars consistently showed the highest rate of canopy accumulation, followed by GR cultivars, while Pre-GR genotypes exhibited the slowest increases. BLUEs, which integrate both environments, reinforced this pattern (Figure 2A). Similar pattern was observed for GY where modern cultivars showed greater average yield (E1:1.5, E2:1.8 kg m^−2^), followed by GR cultivars (E1:1.3, E2:1.6 Kg m^−2^) and cultivars Pre-GR showed lowest GY (E1:1.2, E2:1.2 Kg m^−2^) (Figure 2B).

**Figure 2.**
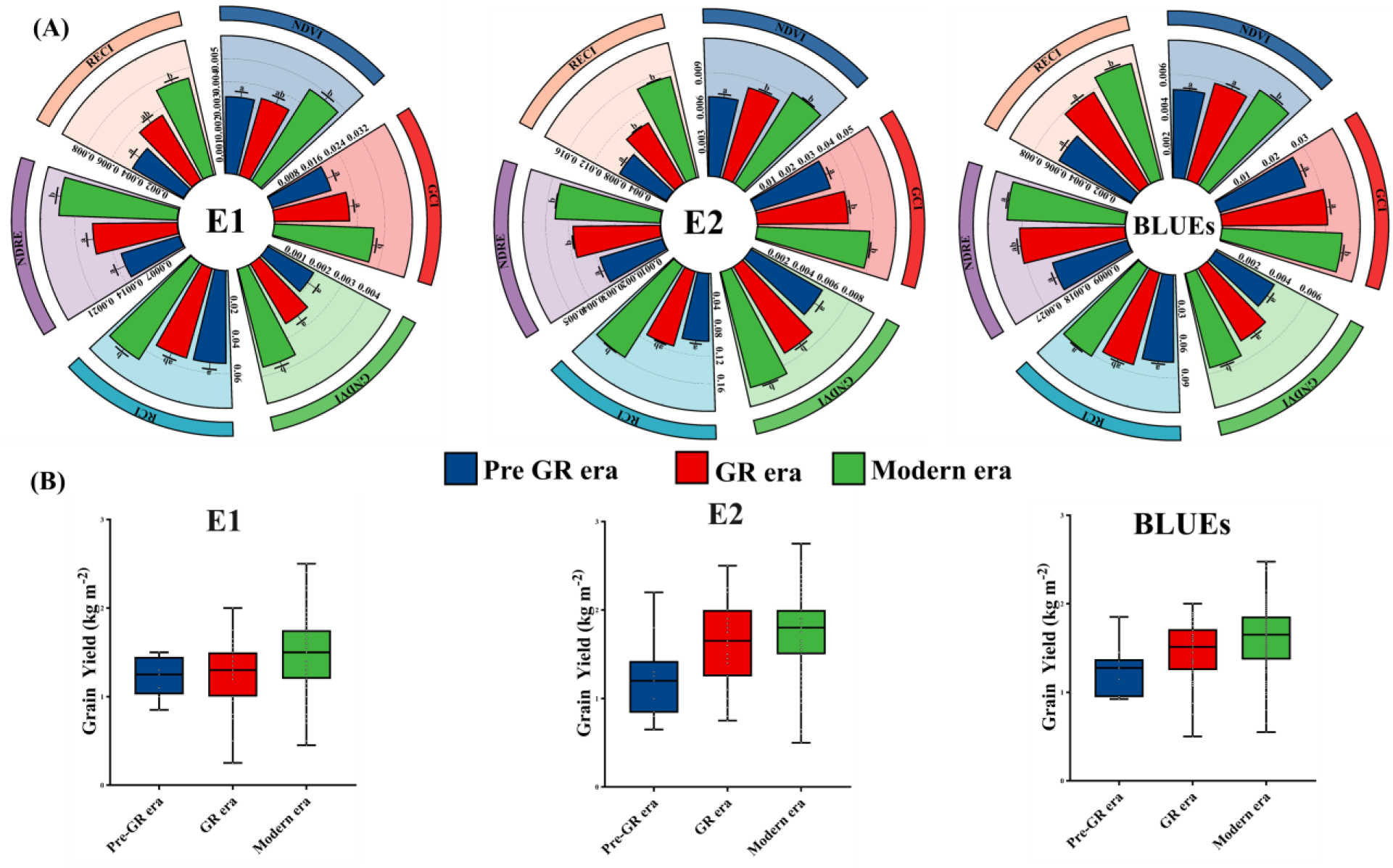
**(A)** Fan plots showing comparison of the six UAV-derived RVIs across three historical breeding eras Pre-GR era, GR era, and Modern era, analyzed in environments E1, E2 and BLUEs. **(B)** Box plot showing Yield trend across three breeding eras in Both environment E1, E2 and their BLUEs.

### 3.2 Correlation and principal component analysis

In E1, all six RVIs showed high correlation between each other (r = 0.845–0.9; p<0.001), while associations with yield were moderate (r=0.28-0.36; p<0.001). GNDVI, GCI showed highest correlation (0.36; p<0.001) while RCI showed lowest (0.28; p<0.001) and TGW moderately related to indices (r = 0.17-0.30; p<0.05-0.001). A similar pattern was also observed in E2, where correlations between RVIs from different VIs remained high (r = 0.78–0.98; p<0.001), while correlations between RVIs and GY were ranged from r= 0.29 to 0.43; p<0.001, and correlations between RVIs with TGW remained relatively low (r = 0.09-0.24; p>0.05-p<0.001). Correlations estimated using BLUEs to capture genetic effects across environments revealed strong associations among the RVIs (r = 0.80-0.9; p<0.001). Their relationships with GY) were moderate (r = 0.37-0.48; p<0.001), while correlations with TGW were relatively stronger than those observed within individual environments (r = 0.21-0.40; p<0.001) (Figure 3).

**Figure 3.**
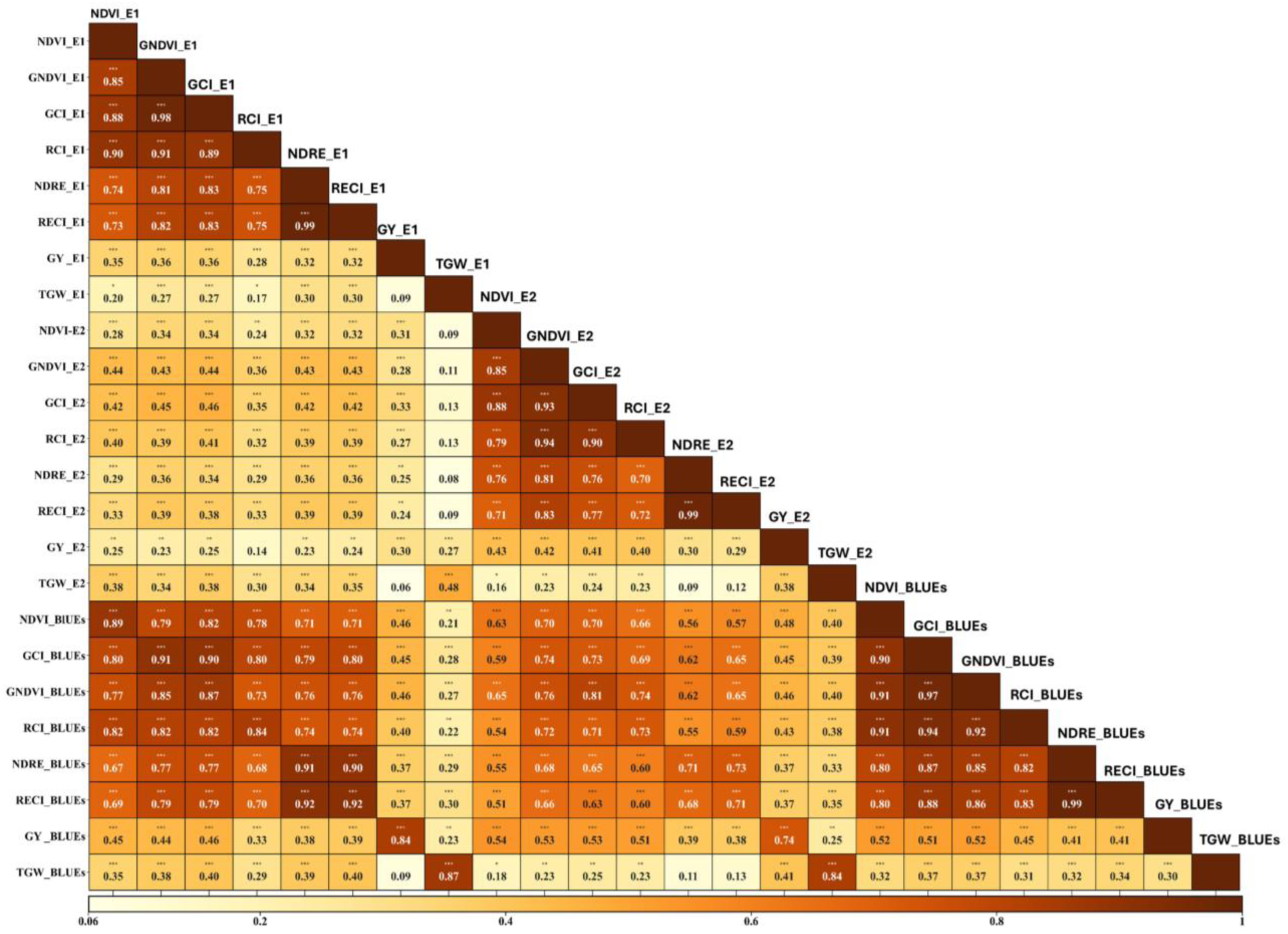
Pearson correlation coefficients among six RVIs and two agronomic traits (GY, TGW) measured across environment E1, environment E2, and BLUEs, showing within-environment consistency and cross-environment relationships. Significance asterisk, P< 0.05=*, P<0.01=**, P<0.001=***.

To evaluate the consistency of RVIs across environments, pairwise correlations were computed among E1, E2, and BLUE values for all RVIs. Across the two years, RVIs exhibited moderate correlations (r = 0.28–0.46; p<0.001), whereas correlations between BLUE values and individual environments were substantially higher (r = 0.63–0.90; p<0.001), indicating improved stability of BLUE-based estimate (Figure 3).

The first two principal components explained a substantial proportion of the total variation (84.4%), with PC1 accounting for 73.2% and PC2 explaining 11.2% (Figure 4). PC1 was largely driven by RVIs, in contrast, PC2 was primarily associated with particularly GY and TGW. In the biplot, there is clear separation for three breeding groups with modern cultivars forming clusters on positive side of PC showing relatively higher RVIs and minimal overlap with cultivars from other two groups, whereas cultivar for Pre-GR era and GR era were more present on negative side of PC1 showing lower RVIs.

**Figure 4.**
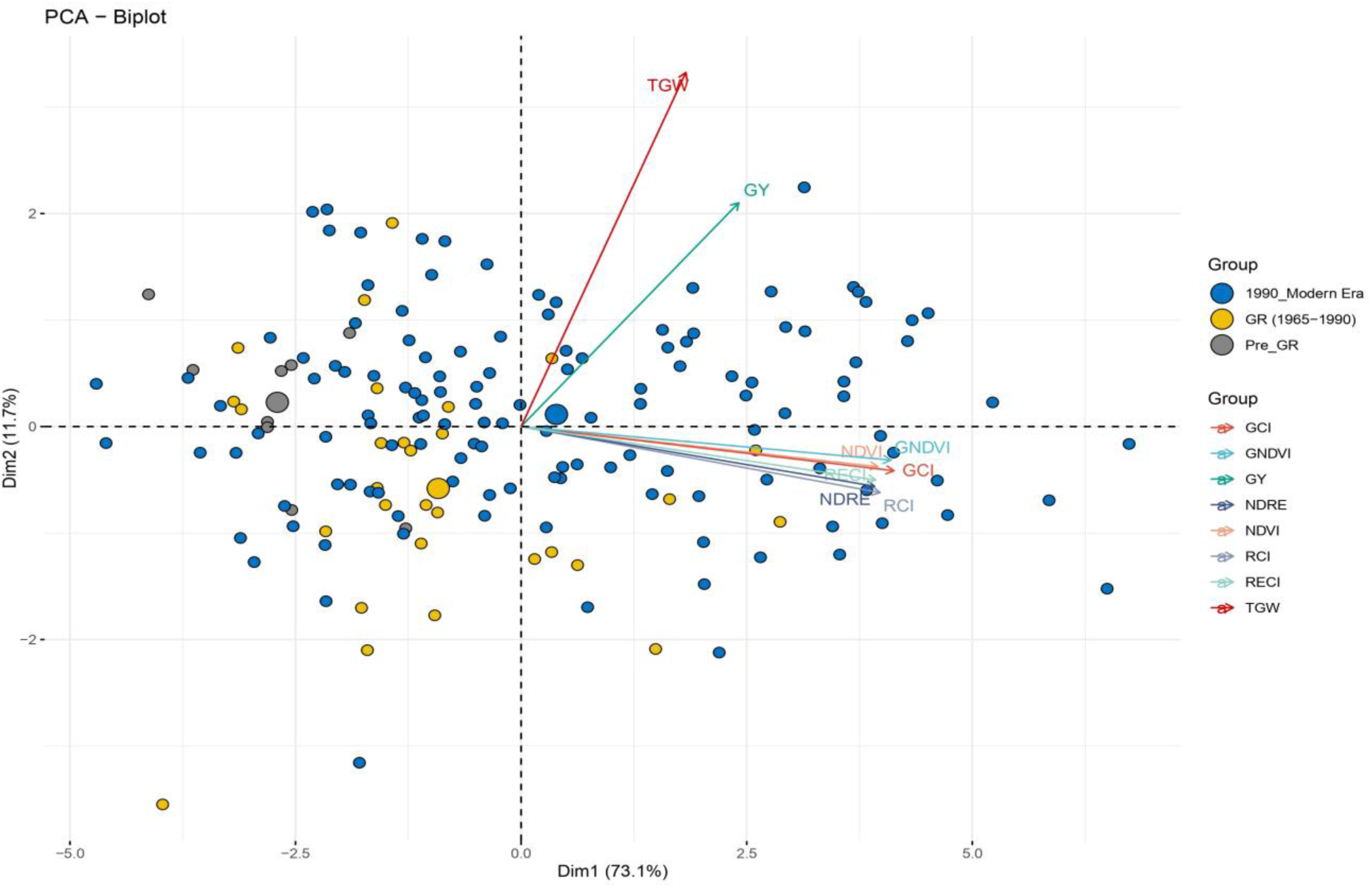
PCA biplot showing the relationships RVIs, GY and TGW, across genotypes from different breeding eras. The first two principal components explain 84.4% of the total variation and points represent genotypes colored by breeding eras. The vectors represent traits.

### 3.3 Genome-wide association analysis

A GWAS was conducted using traits measured in individual environments (E1 and E2) as well as their BLUEs. In total, 328 significant quantitative trait nucleotides (QTNs) were identified, corresponding to 67 unique loci (Supplementary Table S2). Among these, 33 loci exhibited pleiotropic effects, being associated with multiple traits, while the remaining 34 were linked to single traits.

The pleiotropic loci were further classified into two groups (Table 3)(Figure 5). Group I included 12 canopy-specific loci that were associated exclusively with RVIs, with no detectable effects on GY or TGW. Of these, only four loci were consistently detected in more than two environments. Notably, a locus on chr3B at 787 Mb was identified in E2 and BLUEs, while two loci on chr6A (16.57 Mb and 484.41 Mb) were detected only in the BLUEs. The remaining loci in this group were identified in E1 and BLUEs. Importantly, only one stable locus, located on chr5D at ∼76 Mb, was consistently detected across all three environments. These loci which were distributed across multiple chromosomes, including chr1A (581 Mb), chr3B (787 Mb), chr4A (195 and 613 Mb), chr5B (69 Mb), chr5D (76 Mb), and chr6A (611 Mb) exhibited pleiotropic effects across all six RVIs.

**Figure 5.**
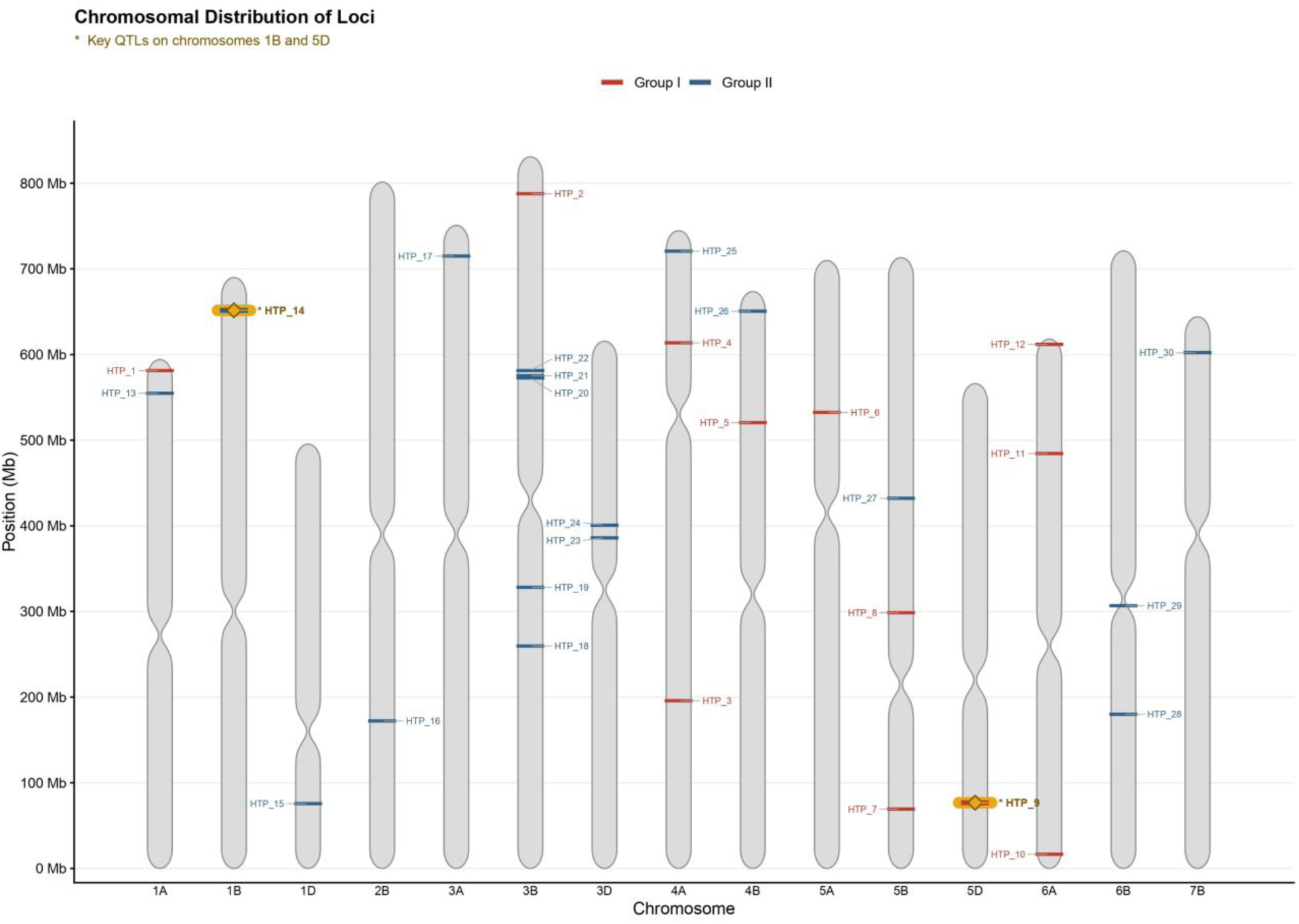
Distribution of pleiotropic loci on chromosomes across two groups, red represents Group I loci (HTP_1 to HTP_12), blue represents Group II loci (HTP_13 to HTP_30). No pleiotropic signals were detected on chr2A, chr2D, chr4D, chr6D, chr7A and chr7D.

**Figure 6.**
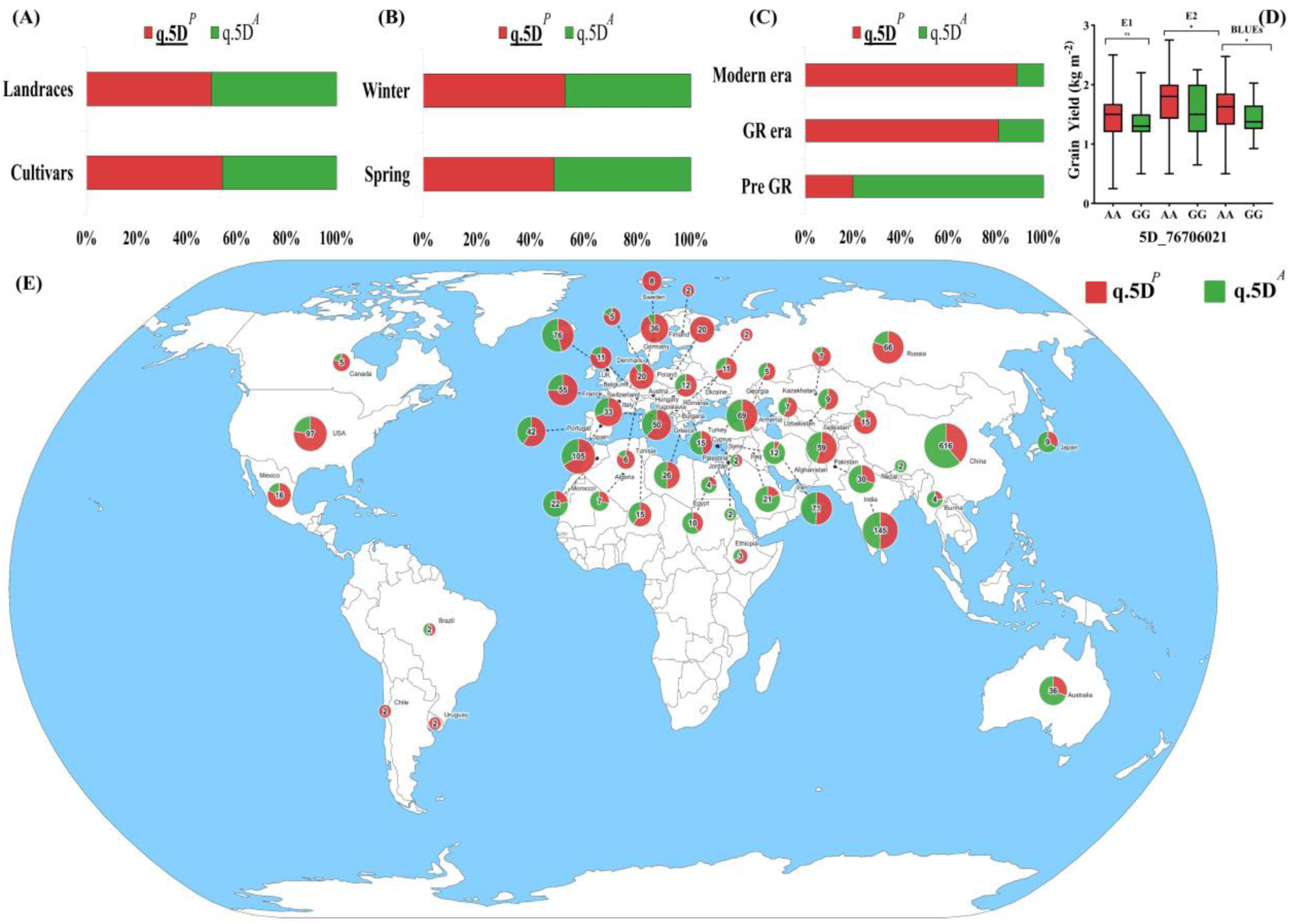
Distribution and allelic effect of the target locus 5D_76706021 (**A)** Distribution of alleles between cultivars and landraces. **(B)** Comparison of allele frequencies between winter-type and spring-type wheat accessions. **(C)** Distribution of alleles in three breeding eras. **(D)** allelic effect of both alleles on the GY in E1, E2 and BLUEs. **(E)** Global distribution of favorable allele. Here *P* represents positive allele and *A* represents alternate allele.

**Table 3.** Summary of significant pleiotropic loci detected through GWAS, categorized into two group according to association pattern.

| Tag SNP<sup>a</sup> | Environment | Traits | Pos/Region (Mb) | P-value Range | MAF | Group |
| --- | --- | --- | --- | --- | --- | --- |
| 1A_581255854 | E1, BLUEs | GCI, GNDVI, NDVI, RCI, RECI, NDRE | 581.26 | 5.52E-05 | 0.25 | Group I |
| 3B_787809760 | E2, BLUEs | GCI, GNDVI, NDVI, RCI, RECI, NDRE | 787.81 | 9.29E-04 | 0.28 | Group I |
| 4A_195803015 | BLUEs | GCI, GNDVI, NDVI, RCI, RECI, NDRE | 195.80 | 1.77E-04 | 0.16 | Group I |
| 4A_613669659 | E1, BLUEs | GCI, GNDVI, NDVI, RCI, RECI, NDRE | 613.67 | 8.89E-05 | 0.31 | Group I |
| 4B_520509762 | E1, BLUEs | GCI, GNDVI | 520.51 | 8.63E-04 | 0.07 | Group I |
| 5A_532472664 | E1, BLUEs | NDVI, RCI | 532.47 | 8.09E-04 | 0.26 | Group I |
| 5B_69305635 | E1, BLUEs | GCI, GNDVI, NDVI, RCI, RECI, NDRE | 69.31 | 9.01E-05 | 0.09 | Group I |
| 5B_298553980 | E1, BLUEs | GCI, GNDVI, RCI, RECI, NDRE | 298-323 | 2.27E-04-8.0E-5 | 0.09 | Group I |
| 5D_76706021 | E1, E2, BLUEs | GCI, GNDVI, NDVI, RCI, RECI, NDRE | 76.71 | 8.15E-04 | 0.17 | Group I |
| 6A_16574141 | BLUEs | RECI, NDRE | 16.57 | 2.11E-04 | 0.12 | Group I |
| 6A_484412806 | BLUEs | RECI, NDRE | 484.41 | 8.16E-04 | 0.23 | Group I |
| 6A_611834945 | E1, BLUEs | GCI, GNDVI, NDVI, RCI,<br>RECI, NDRE | 611.83 | 7.27E-04 | 0.33 | Group I |
| 1A_554807807 | E1, BLUEs | GCI, GNDVI, NDVI, RCI,<br>RECI, NDRE, TGW, GY | 554.81 | 7.06E-04 | 0.12 | Group II |
| 1B_651567054 | E1, E2,<br>BLUEs | GCI, GNDVI, NDVI, RCI,<br>RECI, NDRE, TGW, GY | 624-651 | 2.65E-04-8.47E-05 | 0.38 | Group II |
| 1D_75662834 | E1, E2,<br>BLUEs | GCI, GNDVI, NDVI, RCI,<br>RECI, NDRE, TGW | 75.66 | 7.76E-04 | 0.08 | Group II |
| 2B_172171110 | E1, BLUE | GCI, GNDVI, NDVI, RCI,<br>RECI, NDRE, GY | 172.17 | 7.53E-05 | 0.41 | Group II |
| 3A_714940445 | E1, E2,<br>BLUEs | GNDVI, GY, TGW | 714-747 | 1.08E-04-3.58E-05 | 0.35 | Group II |
| 3B_259661900 | E1, BLUEs | GNDVI, NDVI, TGW | 259.66 | 1.41E-04-2.23E-05 | 0.23 | Group II |
| 3B_328200643 | E1, BLUEs | GCI, TGW | 328.20 | 7.18E-05 | 0.05 | Group II |
| 3B_572684804 | E1, E2,<br>BLUEs | GCI, GNDVI, NDVI, RCI,<br>RECI, NDRE, GY | 572.68 | 9.74E-04 | 0.09 | Group II |
| 3B_575234379 | E1, BLUEs | GCI, GNDVI, NDVI, RCI,<br>RECI, NDRE, GY | 575.23 | 2.13E-05 | 0.28 | Group II |
| 3B_581264249 | E1, E2,<br>BLUEs | GCI, GNDVI, NDVI, RCI,<br>RECI, NDRE, TGW, GY | 581-647 | 1.94E-04-9.3E-06 | 0.35 | Group II |
| 3D_385938484 | E1, E2,<br>BLUEs | GCI, GNDVI, GY, TGW | 10-385 | 1.33E-04-9.98E-05 | 0.13 | Group II |
| 3D_400699255 | BLUEs | NDVI, GY | 400.70 | 8.36E-04 | 0.28 | Group II |
| 4A_720722565 | E2, BLUEs | RECI, NDRE, GY | 713.51-720 | 6.64E-04-1.81E-06 | 0.08 | Group II |
| 4B_650595962 | E1, E2,<br>BLUEs | GCI, GNDVI, NDVI, RCI,<br>RECI, NDRE, TGW | 650-658 | 1.00e-04-2.04E-05 | 0.36 | Group II |
| 5B_432152467 | E1, E2,<br>BLUEs | RECI, NDRE, GY, TGW | 417-654 | 3.73E-04-7.33E-05 | 0.12 | Group II |
| 6B_179999410 | E2, BLUEs | NDVI, GY | 171-205 | 3.1E-04-2.54E-05 | 0.11 | Group II |
| 6B_306776154 | E1, BLUEs | GCI, GNDVI, TGW | 306.78 | 8.69E-06 | 0.10 | Group II |
| 7B_602265277 | E1, E2,<br>BLUEs | GCI, GNDVI, NDVI, RCI,<br>RECI, NDRE, TGW, GY | 158-605 | 1.5E-04-9.98E-05 | 0.17 | Group II |

<sup>a</sup> SNP name represents chromosome number followed by SNP position

Group II consisted of 18 multi-trait loci simultaneously associated with GY, TGW and at least one RVI. Out of these 18 loci, 9 stable loci were identified across all environments (E1, E2and BLUEs). A total of 6 loci showed pleiotropic association between RVIs and GY. Whereas only 5 loci were pleiotropic for RVIs and TGW. Remaining 7 loci were pleiotropic for both GY and TGW along with RVIs. loci on chr1A (554 Mb), chr1B (624 Mb), chr3B (581 Mb), and chr4B (650 Mb) were associated with all RVIs as well as GY and TGW. A stable locus on chr3A (714 Mb) showed associations of GNDVI with both GY and TGW, while a locus on chr3D (10 Mb) was associated with GCI and GNDVI alongside yield traits. Additionally, a locus on chr4A (713 Mb) was associated with RECI and NDRE together with GY. A stable Locus on chr3B∼572 Mb was associated with all RVIs and GY.

The three pleiotropic loci on chr3B showed associations exclusively between GY and TGW without RVI involvement and were not assigned to either group.

### 3.4 Allelic distribution and effects of key loci

Two stable loci on chr5D and chr1B were further characterized for their allelic effects and frequencies in global wheat collections. Allelic effect analysis for chr5D showed that allele (AA) was consistently associated with higher GY (E1-1.43 kg m^−2^and E2-1.72 kg m^−2^) than allele (GG) (E1-1.35 kg m^−2^ and E2-1.53 kg m^−2^) across all three environments (Figure 7). The favorable allele of 5D was present in ∼20% Pre-Green Revolution cultivars and its frequency constituently increased to 80% in the GR cultivars and reached ∼90% in modern cultivars. In global wheat collection of ∼3000 cultivars, the frequency of favorable allele was approximately 50% in landraces and 55% in cultivars, similarly in terms of growth habitat favorable allele was present more in winter type 55% than spring type ∼50%.

**Figure 7.**
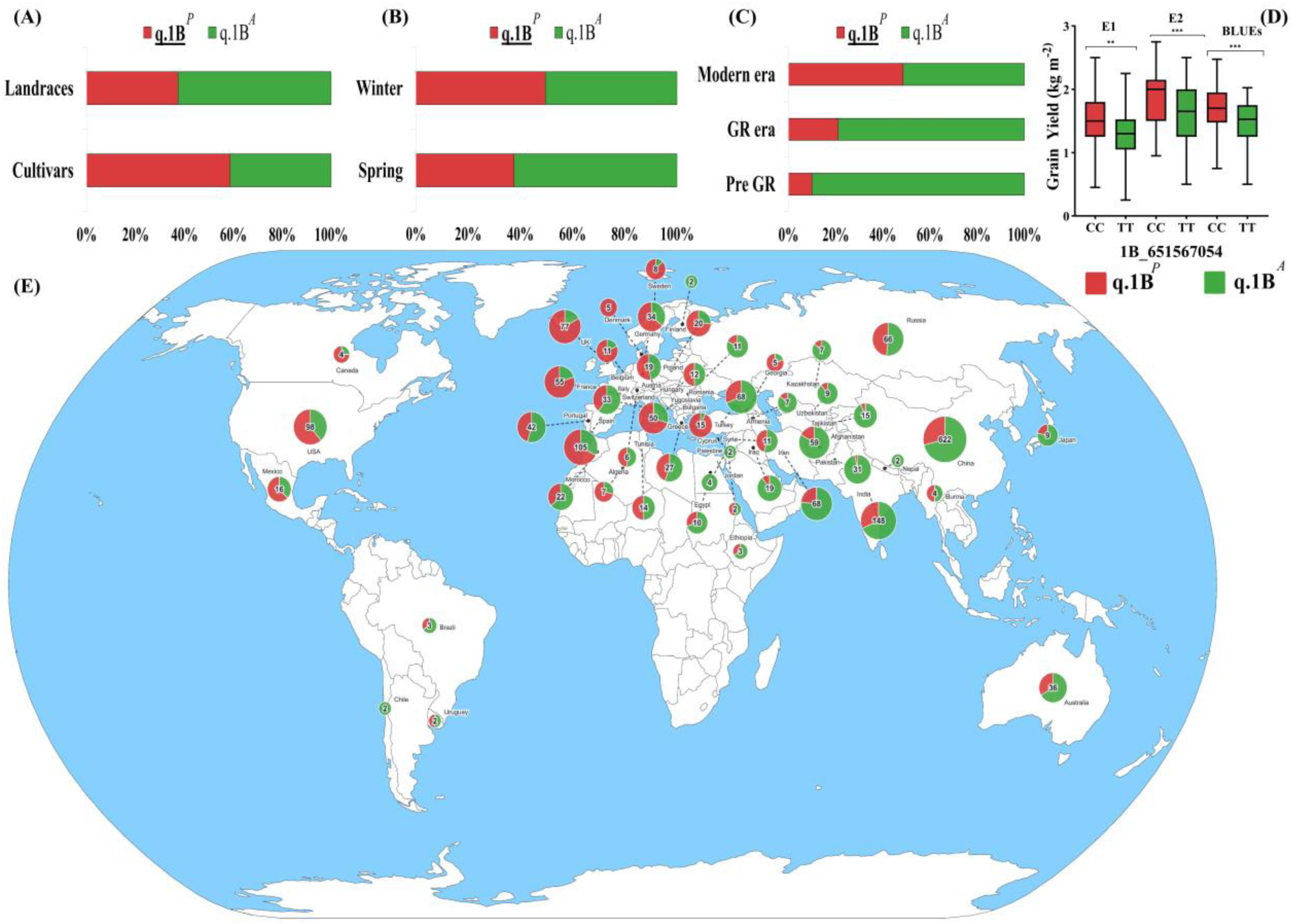
Distribution and allelic effect of the target locus1B_651567054 (**A)** Distribution of alleles between cultivars and landraces. **(B)** Comparison of allele frequencies between winter-type and spring-type wheat accessions. **(C)** Distribution of alleles in three breeding eras. **(D)** allelic effect of both alleles on the GY in E1, E2 and BLUEs. **(E)** Global distribution of favorable allele. Here *P* represents positive allele and *A* represents alternate allele.

For chr1B locus, allelic distribution showed that allele (CC) was consistently associated with higher GY (E1-1.52 kg m^−2^ and E2-1.86 kg m^−2^) than allele (TT) (E1-1.34 kg m^−2^ and 1.60 kg m^−2^). The favorable allele was present in ∼10 % Pre-Green Revolution Cultivars and its frequency increased to ∼25% in GR cultivars and reached ∼50% in modern cultivars. In global wheat collection, the frequency of favorable allele was 40% in landraces and 60% in cultivars, similarly across growth habitat favorable allele was present more in winter type ∼50% than spring type 40% (Figure 7).

### 3.5 Development of KASP assays

KASP markers developed for two key loci were tested in a population of 191 Chinese wheat genotypes (Figure 8). KASP markers showed clear separation of alleles forming clear separate clusters for each allele.

**Figure 8.**
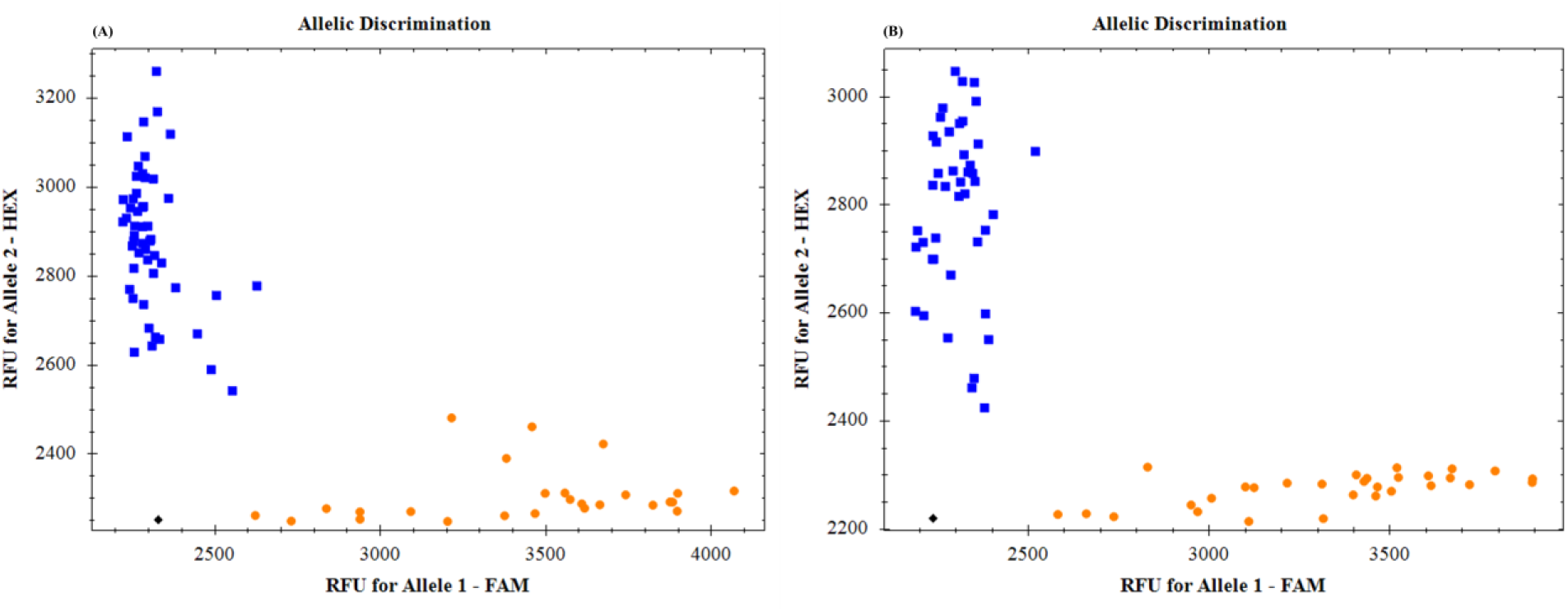
KASP genotyping scatter plots for the 5D_76706021(A) and 1B_651567054 (B). The x-axis represents the fluorescence intensity for the FAM allele, and the y-axis represents the fluorescence intensity for the HEX allele. Blue boxes denote HEX fluorescence clusters (allele 2), and yellow dots denote FAM fluorescence clusters (allele 1), illustrating clear separation of genotypic classes.

### 3.6 Potential candidate gene near Key loci

To identify genes potentially underlying the key loci representing both groups, the associated genomic intervals were examined using the wheatomics resource for gene annotation and expression profiling.

For Group-II key loci four candidate gene were identified. TraesCS1B01G418400 (642.39–642.39Mb; AUX/IAA domain, PF02309), TraesCS1B01G424100 (647.22–647.23Mb; Homeobox domain, PF00046), TraesCS1B01G424700 (649.04–649.06 Mb; Auxin-responsive domain, PF06507, B3 DNA binding domain PF02362), and TraesCS1B01G425200 (649.67–649.70 Mb; Glutaredoxin domain, PF00462).

For Group I key loci on chr5D two putative candidate genes were identified. TraesCS5D01G076400 (Glycosyltransferase domain; PF00201) and TraesCS5D01G076800 (B3 DNA-binding domain; PF02362).

Expression profiles from the wheat omics database indicated that these genes are actively expressed in root, shoot and developing spike tissues during vegetative and early reproductive stages, corresponding to the developmental window evaluated in the GWAS (Supplementary Figure S1).

## 4. Discussions

### 4.1 UAV-based RVIs capture pre-heading biomass accumulation for grain formation

In this study, UAV-derived rates of RVIs were successfully used to quantify canopy and biomass development during the vegetative stage, providing a dynamic proxy for early biomass accumulation in wheat. The strong variation in RVIs observed across environments, together with consistent ranking among breeding groups, indicates that these traits effectively capture genotypic differences in canopy development despite environmental influence. Rapid canopy expansion during this stage is critical, as it governs light interception, radiation use efficiency, and the establishment of photosynthetic capacity that ultimately supports reproductive development and grain filling. Previous studies have demonstrated that early canopy closure and efficient radiation interception during vegetative growth strongly influence crop growth rate and final yield potential (Aisawi et al., 2015; Foulkes et al., 2011).

A key limitation in wheat improvement is the near saturation of harvest index, beyond which further gains may compromise canopy photosynthetic areas, stem carbohydrate reserves, and structural integrity. The observed higher RVIs and grain yield in modern cultivars across environments suggest that breeding progress has increasingly relied on improved canopy development and biomass accumulation rather than further increases in harvest index. This aligns with evidence that future yield gains must be achieved through enhanced total biomass production, supported by improved radiation use efficiency and canopy light interception, traits largely determined during the pre-heading vegetative phase (Fischer et al., 2024; Foulkes et al., 2022; Rivera-Amado et al., 2019).

By integrating multiple UAV timepoints, RVIs capture the rate of canopy establishment, providing an effective measure of the metabolic momentum of biomass accumulation that is not accessible through conventional phenotyping approaches (Song et al., 2025). Within the source–sink framework, the canopy acts as the primary assimilate source, while developing spikes become dominant sinks after anthesis(Reynolds et al., 2022). Genotypes with higher RVIs establish a stronger source canopy before anthesis, which is reflected in their higher yield performance.

The moderate correlations observed between RVIs and grain yield (r = 0.28–0.51) are consistent with this upstream role of canopy development, as final yield is also influenced by post-heading processes such as grain filling efficiency, assimilate remobilization, and stress responses. The weaker correlations between RVIs and TGW further support this interpretation, indicating that RVIs primarily capture variation in pre-heading source establishment rather than sink-driven grain filling processes (Foulkes et al., 2022).

These findings are consistent with previous studies showing that VIs tend to exhibit stronger relationships with yield when measured during grain filling (Hassan et al., 2019), whereas vegetative-stage measurements capture earlier physiological processes contributing to yield formation. Collectively, the results demonstrate that UAV-based RVIs provide a robust and biologically meaningful approach to quantify early canopy development, enabling improved understanding of biomass accumulation and its contribution to yield potential in wheat.

### 4.2 Historical breeding trends in pre-heading canopy development along with yield

Results showed progressive increase in RVIs from Pre-Green Revolution to Green Revolution and modern cultivars, observed consistently across contrasting environments and reinforced by BLUEs. Modern cultivars exhibited the highest rates of pre-heading canopy development, followed by Green Revolution-era materials, while Pre-Green Revolution genotypes showed the slowest canopy expansion rates. This directional trend closely paralleled improvements in GY, with modern cultivars outperforming earlier groups across environments. This pattern is likely attributable to the accelerated rate of RVI accumulation observed in modern genotypes. Principal component analysis further supported this relationship, with RVIs loading strongly on PC1 and yield traits (GY and TGW) on PC2, indicating that canopy development rate represents a distinct biological dimension rather than reflecting yield outcomes. Importantly, these results demonstrate that UAV-based RVIs can successfully monitor historical breeding progress in canopy development and biomass accumulation. Although breeding during the Green Revolution primarily targeted yield, harvest index, and lodging resistance, it also indirectly improved early canopy establishment, likely through enhanced photosynthetic capacity and nitrogen-use efficiency. This interpretation aligns with previous reports of genetic gains in early vigor, radiation-use efficiency, and biomass production in modern wheat germplasm (R. A. Fischer & Edmeades, 2010; Sadras & Lawson, 2011). The ability of UAV-derived RVIs to capture these trends highlights their strong potential as scalable, high-throughput phenotyping tools. Unlike conventional methods, UAV platforms enable rapid, non-destructive, and repeated measurements across large field trials, making RVIs particularly suitable for integration into large breeding programs. Therefore, RVIs not only reflect past genetic gains but also offer a practical and efficient trait for direct selection, providing a pathway to accelerate improvement in pre-heading canopy development and overall yield potential in future wheat breeding efforts.

### 4.3 Genetic architecture of pre-heading canopy development and yield formation

Conventional GWAS typically analyzes yield as a final outcome trait and therefore identify statistical associations at the system output level, often without clarifying the pre-anthesis biological mechanisms that contribute to yield formation. The pathway from genotype to yield involves a sequence of intermediate developmental processes, each under partial genetic control, whose individual contributions are masked when collapsed into a single end-season measurement limiting mechanistic interpretation. The identification of Group I loci exclusively associated with RVIs, but not with GY or TGW, suggests that UAV-derived canopy traits can reveal genomic regions controlling pre-heading source establishment that may be missed by conventional yield-based GWAS. These loci likely regulate upstream vegetative processes such as canopy expansion, biomass accumulation, chlorophyll development, and radiation interception before yield components are fully determined. This interpretation is consistent with previous studies showing that UAV- and ground-based VIs can capture genetic variation in canopy vigor, NDVI, chlorophyll content, and adaptive physiological traits in wheat (Cheng et al., 2025; Condorelli et al., 2018; Rutkoski et al., 2016). The broad-spectrum Group I loci associated with all six RVIs, including regions on chr1A, chr3B, chr4A, chr5B, chr5D, and chr6A, may represent regulatory hubs influencing general canopy development. Their effects across multiple indices suggest coordinated control of canopy closure, biomass accumulation, and photosynthetic surface area. In contrast, index-specific loci provide greater physiological resolution. For example, loci associated with GCI and GNDVI likely reflect variation in canopy greenness and chlorophyll concentration, while NDVI and RCI-associated loci may be more closely related to canopy structure, biomass, and radiation interception. Similarly, loci associated with NDRE and RECI indicate genetic regulation of red-edge reflectance, which is commonly linked with chlorophyll concentration, nitrogen status, and photosynthetic capacity (Delegido et al., 2011; Fitzgerald et al., 2010; Hatfield et al., 2008). This supports the value of using multiple RVIs rather than a single index, because different spectral indices can partition canopy development into structural, pigment-related, and nitrogen-related components. Group II loci, which were associated with both RVIs and yield-related traits, represents genomic regions where pre-heading source development converges with final yield formation. loci associated with all six RVIs together with GY and TGW may reflect integrative source–sink regulators whose effects on early canopy development are translated into yield advantages. The consistent detection of loci on chr1B, chr3B, and chr4B across environments further strengthens their potential utility for breeding. Previous studies have similarly shown that spectral indices derived from high-throughput phenotyping platforms can improve prediction of grain yield and identify genomic regions linked to yield-related physiological traits in wheat (Condorelli et al., 2018; Mérida-García et al., 2024; Rutkoski et al., 2016; Sharma et al., 2024). The loci associated with RVIs and GY but not TGW are particularly informative, as they suggest that early canopy development may influence yield mainly through grain number rather than individual grain weight. This agrees with wheat source–sink physiology, where assimilate availability before anthesis supports spike growth, floret survival, and grain number determination (Foulkes et al., 2011; Gonzalez-Navarro et al., 2016; Reynolds et al., 2022). In contrast, TGW is more strongly governed by post-anthesis grain filling, endosperm development, starch accumulation, and remobilization of stored assimilates.

Overall, these results show that UAV-based multi-index GWAS can dissect both independent source-establishment loci and source–sink convergence loci. Group I loci provide access to vegetative canopy-development genetics that are not directly reflected in final yield, whereas Group II loci identify genomic regions where early canopy vigor contributes to yield formation.

### 4.4 Directional selection and breeding implications

Allele frequency changes at loci chr5D and chr1B reveal clear signs of productivity driven selection during wheat improvement. At chr5D, the favorable AA allele increased from low frequency in pre–Green Revolution germplasm to near fixation in modern cultivars, a directional shift consistent with strong indirect selection as a correlated response to yield improvement during modern breeding, possibly through its effects on canopy development and vigor captured by RVIs. Its similar frequency across growth habits suggests it provides a canopy establishment advantage that is broadly expressed regardless of season length. The relatively small increase in favorable allele frequency from landraces to cultivars indicates that this allele may already have been present at moderate frequency in traditional germplasm and was subsequently enriched during modern breeding.

At chr1B, the favorable CC allele showed a more gradual but steady increase from landraces to modern cultivars, reflecting its progressive incorporation under formal high-input breeding conditions where its direct contribution to yield components is consistently expressed. Its higher frequency in winter compared with spring types suggests that this locus may influence canopy development traits that are more strongly expressed under longer growing seasons. Thus, although both loci contribute to yield improvement, their increasing frequencies across breeding history likely reflect different selection dynamics.

Importantly, favorable allele CC at 1B has not reached complete fixation, which is still present in only ∼50% of modern cultivars, indicating remaining potential for genetic gain through targeted marker-assisted selection within elite germplasm. Overall, combining allele frequency trends with GWAS-based effect estimates provides genomic evidence that historical yield gains in Pakistani wheat have been accompanied by the gradual enrichment of loci associated with canopy efficiency, highlighting both regions as practical targets for accelerating future improvement.

The KASP markers developed for the two stable loci showed high call rates and clear allele discrimination in an independent wheat panel, demonstrating reliable assay performance. The successful conversion of peak SNPs into KASP assays highlights their suitability for high-throughput genotyping and potential application in marker-assisted selection. These markers provide practical tools for screening breeding germplasm and tracking favorable alleles in wheat improvement programs. Further validation in diverse populations and environments may strengthen their utility for breeding applications.

### 4.5 Candidate gene identification

The candidate genes at the chromosome 1B locus (Group II) suggest that this region regulates growth processes extending beyond vegetative development into reproductive performance. TraesCS1B01G418400 contains an AUX/IAA domain (PF02309) characteristic of Aux/IAA repressors involved in auxin-dependent transcriptional control of tiller development and organ differentiation (Reed, 2001), while TraesCS1B01G424700 encodes a B3 DNA-binding domain (PF02362) representative of ARF-type transcription factors that act as the activating counterpart in auxin signaling during organ initiation and developmental transitions (Guilfoyle & Hagen, 2007). The co-occurrence of both repressor and activator components within the same locus points to coordinated auxin-responsive transcriptional regulation. TraesCS1B01G424100 carries a Homeobox domain (PF00046) associated with meristem identity and developmental phase transitions that coordinate vegetative architecture with spike development in cereals (Mukherjee et al., 2009), and TraesCS1B01G425200 encodes a Glutaredoxin domain (PF00462) linked to redox regulation of cell proliferation under developmental cues (Rouhier et al., 2008). This combination of hormone signalling, transcriptional regulation, and redox control provides a mechanistic basis for the association of this locus with both composite growth indices and final grain yield.

In contrast, the chr 5D locus (Group I) harbors genes whose functions are primarily associated with vegetative growth. TraesCS5D01G076400 possesses a glycosyltransferase domain (PF00201) central to cell wall biosynthesis and cell expansion (Lairson et al., 2008), while TraesCS5D01G076800 contains a B3 DNA-binding domain (PF02362) characteristic of plant transcription factors involved in hormone-mediated developmental responses (Swaminathan et al., 2008). The absence of reproductive-stage functions within this locus is consistent with its exclusive association with VIs. The expression of these candidate genes during vegetative and early reproductive stages aligns with the developmental window captured by the rate-of-gain indices, supporting their biological relevance to the pleiotropic signals detected in GWAS.

### 4.6 RVIs as a window into phenotypic black box of grain yield

GY represents a single terminal number that integrates a tightly coordinated developmental sequence in which canopy source capacity is first established and then mobilized into sink formation. When only this endpoint is mapped, the individual genetic contributions of the intermediate developmental processes that generate it are systematically diluted or invisible. Conventional GWAS operates on this output, capturing what the plant produced rather than how it produced it, leaving the regulatory architecture of the pre-heading developmental phase effectively as a phenotypic black box.

RVIs address this directly by providing the scalable phenotypic access to the pre-heading phase. The Group I loci identified without yield association represents genetic control of processes occurring predominantly within this black box real biological regulation that conventional yield mapping would not effectively detect. Group II loci, by connecting the source developmental phase to final yield, reveal where the black box opens into the yield formation pathway. Together, these loci do not merely confirm yield-associated genomic regions. They begin to map the biological pathway and architecture that generates yield. In this sense, RVIs help bridge the phenotypic gap in the pre-heading phase, improving understanding of the genetic basis of yield formation.

### 4.7 Conclusion and prospects

The present study supports a time-series data acquisition which captures the development phase to understand complex traits like grain yield. UAV-derived vegetation index accumulation rates from tillering to heading effectively captured heritable variation in pre-heading canopy development and enabled the identification of 67 genomic regions across all 21 wheat chromosomes through GWAS. These dynamic RVIs also revealed historical breeding progress, with modern cultivars consistently exhibiting faster canopy development than Green Revolution and pre-Green Revolution lines, demonstrating their ability to resolve temporal patterns of genetic gain that static measurements cannot capture. The classification of pleiotropic loci into canopy-specific (Group I) and yield-associated (Group II) categories revealed a hierarchical genetic architecture underlying pre-heading source establishment and its linkage to yield formation. Notably, the identification of favorable but unfixed alleles and environmentally stable loci highlights immediate opportunities for targeted genetic improvement. Importantly, the development and validation of KASP markers for selected loci further strengthen the translational value of these findings. These markers provide a practical tool for breeders to efficiently introgress favorable alleles controlling canopy development and biomass accumulation into elite germplasm. Overall, this study establishes UAV-derived RVIs as scalable and biologically meaningful surrogate traits for dissecting early biomass accumulation and its genetic basis. By integrating dynamic UAV phenotyping with genomic tools such as KASP markers, this work provides a powerful framework to uncover and deploy genetic variation that remains inaccessible to conventional endpoint measurements, thereby accelerating genetic gain and improving yield potential in wheat breeding programs.

## Acknowledgements

This study was financially supported by Science and Technology Partnership Program (Project No. KY202201001) of Ministry of Science and Technology of the Peoplés Republic of China (MoST, China), Higher Education Commission’s National Research Program for Universities (NRPU 20-15269), and CIMMYT-SDAU joint project.

## Authors contributions

SR performed the core experiments and data analysis; AR (Ali Raza), ZH (Zijian He), and MSA collected the phenotyping data; ZM and MF managed and executed the field trials; AR (Awais Rasheed), MAH, LL, and MW designed the experiments; SR wrote the original draft; and JW, YX, ZH (Zhonghu He), and AR (Awais Rasheed) reviewed and revised the manuscript

## Data availability

All genotypic and phenotypic data are available upon reasonable request.

