## Supplementary Figure S1 for "Discovering genetic loci associated with rate of vegetative index gain using UAV-based phenomics in spring wheat"

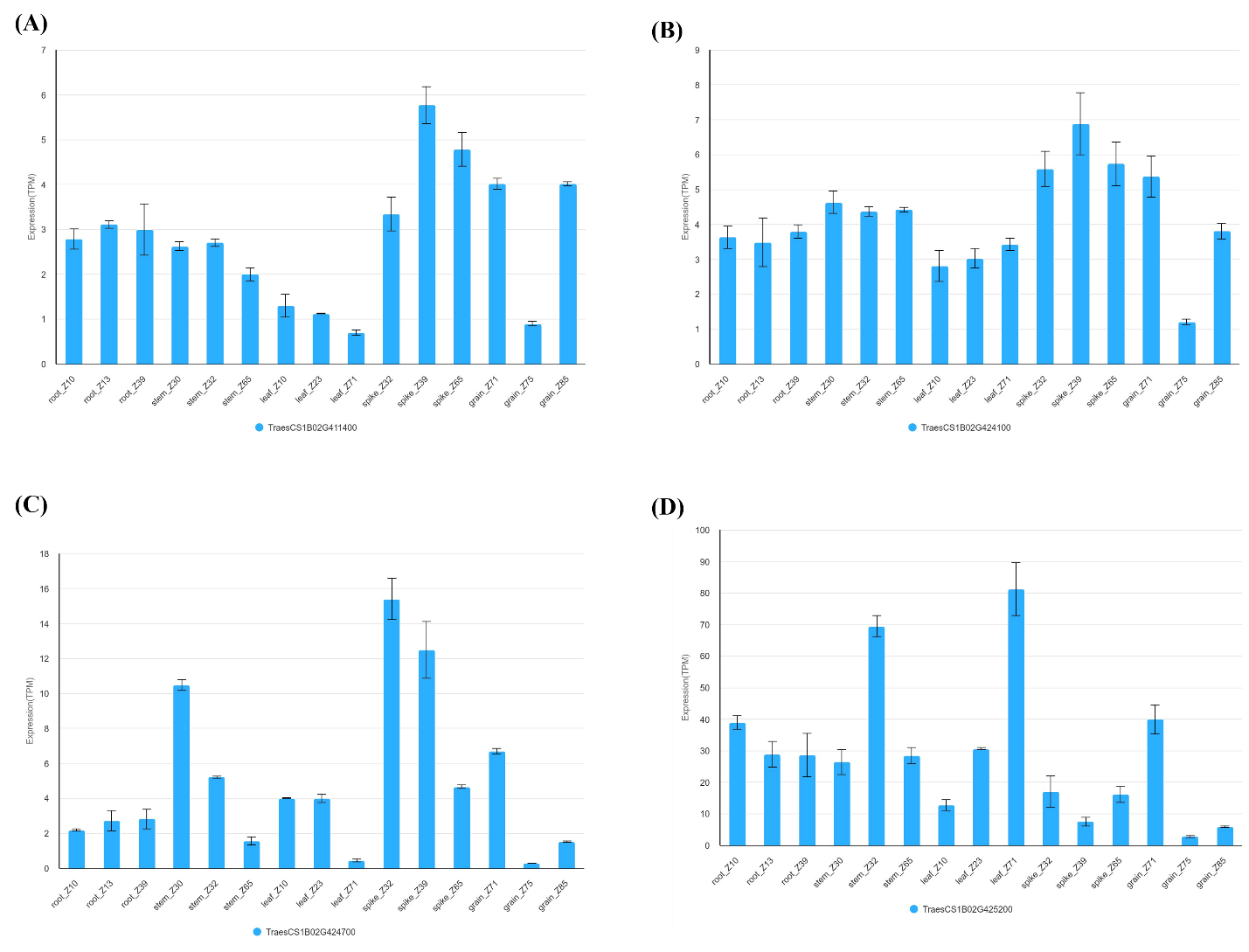

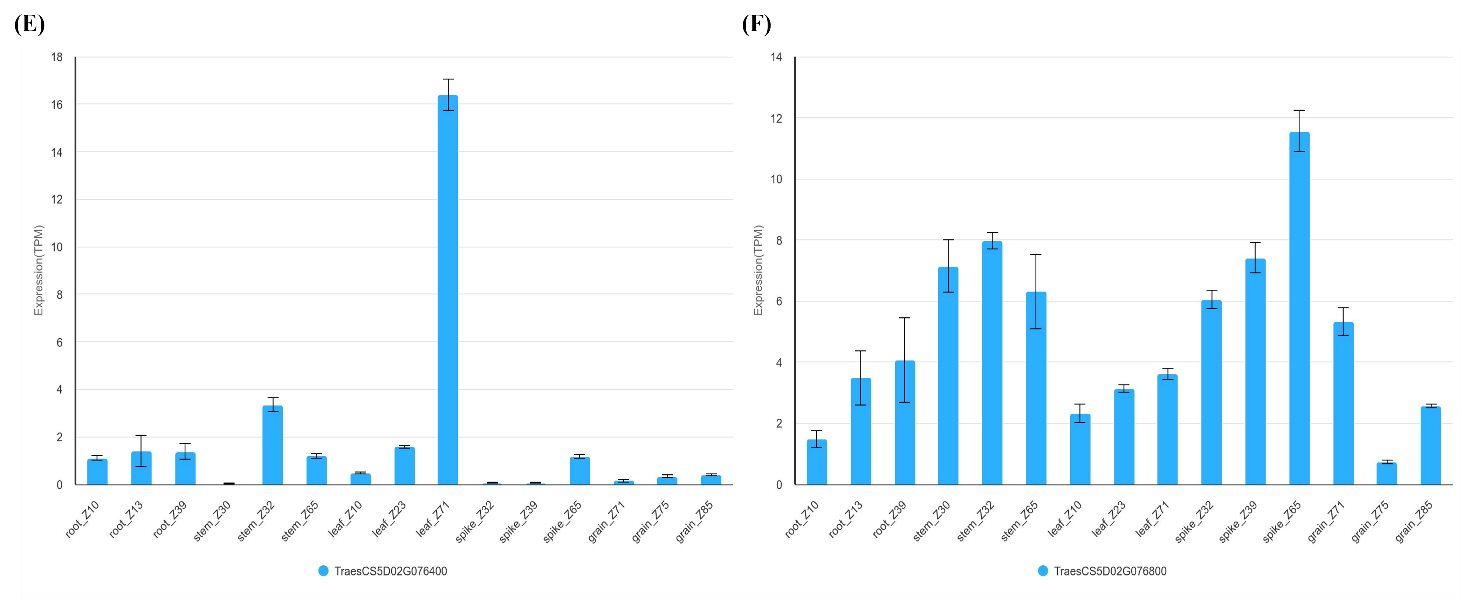

**Supplementary Figure S1:** Expression data bar plot from wheat omic database **(A)** TraesCS1B01G418400 (Auxin-responsive domain), **(B)** TraesCS1B01G424100 (Homeobox domain), **(C)** TraesCS1B01G424700 (Auxin-responsive domain), **(D)** TraesCS1B01G425200 (Glutaredoxin domain), **(E)** TraesCS5D01G076400 (Glycosyltransferase domain; PF00201) and **(F)** TraesCS5D01G076800 (B3 DNA-binding domain; PF02362).
